# A phototaxis assay to measure sublethal effects of pesticides on bees

**DOI:** 10.1101/2025.02.11.637502

**Authors:** Gonzalo Sancho, Sergio Albacete, Celeste Azpiazu, Fabio Sgolastra, Anselm Rodrigo, Jordi Bosch

## Abstract

Pesticides are considered a main driver of world-wide bee declines. In agricultural areas, bees are exposed to combinations of pesticides at low concentrations. The extent to which these low levels may cause sublethal effects remains unknown. Laboratory methods to detect sublethal effects are needed as a first step in assessing the potential hazards of pesticides at low concentrations. Current bee risk assessment schemes rely on a single species, the highly social *Apis mellifera*, and provide insufficient coverage of sublethal effects. Due to fundamental life history differences, available tests cannot be applied to solitary bees. We provide a simple phototaxis assay to detect sublethal pesticide effects on bees. The assay is highly effective (only 6.63% of the bees failed to respond) and provides an unambiguous binary response (bees either walk straight to the light source or walk erratically across the arena). We validate the assay by conducting two experiments. In the first one, we estimate dose-response curves and calculate ED50 and benchmark dose (BMD) values of an insecticide on *Osmia bicornis* and *O. tricornis*. In the second one, we assess the effects of three insecticide doses, alone and in combination with a fungicide, in *O. cornuta* and *A. mellifera*. These experiments show that our assay can detect effects of field-realistic levels of acetamiprid exposure as low as 1-30 ng/bee. The phototaxis assay can be used to obtain relevant ecotoxicological endpoints at low sublethal concentrations in both solitary and honey bees, thus contributing to fill an important gap in bee risk assessment.

## 1. Introduction

Bee abundance and diversity have been declining in recent decades, and pesticide use is considered one of the main drivers of these declines (Dicks et al., 2021). Studies analysing pesticide residues in the pollen and nectar of crop flowers (Azpiazu et al., 2023a; Heller et al., 2020; Zioga et al., 2020), wild flowers (Botías et al., 2015; David et al., 2016; Ward et al., 2022) and bee food provisions (Graham et al., 2021; Nicholson et al., 2023; Porrini et al., 2016) reveal that the presence of combinations of pesticides (including insecticides, fungicides and herbicides) is widespread in agricultural environments. These “pesticide cocktails” are typically found at low concentrations and, for the most part, are not expected to have acute lethal effects on bees (Azpiazu et al., 2023a; Heller et al., 2020; Siviter et al., 2021; Zioga et al., 2020). However, the extent to which these pesticide combinations may have sublethal effects remains largely unknown. Because they do not result in short-term death, sublethal effects are often difficult to quantify or even detect, both in the field and in the laboratory. However, even if less severe than lethal effects, sublethal effects can significantly impact individual fitness, with potential consequences on population dynamics. A study analysing pesticide residues in bumblebee, *Bombus terrestris*, pollen stores, shows that colony growth is negatively affected by pesticide exposure across European landscapes (Nicholson et al., 2023). Another study shows that exposure to sublethal insecticide-fungicide concentrations causes a chain of effects, ultimately resulting in lower female fecundity and negative population growth in the solitary bee *Osmia cornuta* (Albacete et al., 2024).

Given the pervasiveness of low levels of pesticide residues in flowers and bee matrices, reliable methods for detecting sublethal effects in the laboratory are needed to assess the potential hazard of low pesticide and pesticide mixture concentrations on bees. Sublethal tests based on physiological, behavioural, and cognitive endpoints for honey bees (*Apis mellifera*) are well established, standardized and partially included in pesticide risk assessment schemes (EFSA, 2013; USEPA, 2014). These tests include the proboscis extension reflex test (Decourtye et al., 2004), the homing flight test (Henry et al., 2012; OECD, 2021), and the hypopharyngeal gland development test (Hatjina et al., 2013; Renzi et al., 2016). However, due to fundamental differences in life history traits, these tests are not easily applicable to non-social bees. The proboscis extension reflex is readily elicited in social bees and wasps (eg. Takeda, 1961; Laloi *et al*., 1999; Mc Cabe *et al*., 2007; Gong, Tan and Nieh, 2019), but not in solitary bees (Vorel and Pitts-Singer, 2010). Similarly, solitary bees are unlikely to find and use artificial syrup feeders in the field, thus hindering their training and preventing the implementation of the homing flight test. Finally, solitary bees do not produce royal jelly to feed larvae and the ontogeny of their hypopharingeal glands, which are much less developed than those of social bees, is poorly understood (Cruz-Landim, 1967). As a result, standardized sublethal methods for solitary bees are lacking (Tosi et al., 2022). The focus on solitary bees is important for several reasons. First, most bee species worldwide (ca. 90%) are solitary or cleptoparasitic of solitary bees (Danforth et al., 2019). Second, solitary bees differ from honey bees in terms of fundamental behavioral and life history traits, resulting in different routes and levels of exposure (Sgolastra et al., 2019). Third, different bee species show different levels of sensitivity to pesticides. Solitary bees, in particular, have been shown to be more sensitive than honey bees to various insecticides (Arena and Sgolastra, 2014; Azpiazu et al., 2021; Sgolastra et al., 2017; Uhl et al., 2019). For these reasons, the European Food Safety Authority has recommended the use of two solitary bee species (*O. cornuta* and *Osmia bicornis*) as model species in pesticide risk assessment (EFSA, 2013). However, the inclusion of these species in risk assessment schemes is being delayed, partly due to the lack of standardized methodologies.

In recent years, we have developed a simple phototaxis-based laboratory assay to measure sublethal pesticide effects in bees. Phototaxis is the movement of a motile organism in response to light, either towards it (positive phototaxis) or away from it (negative) (Jékely 2009). Positive phototaxis has been used to monitor insect populations (e.g., Jonason, Franzén and Ranius, 2014) and to control pests (e.g., Kim, Huang and Lei, 2019). In bees, phototaxis has been used to study learning and memory, and to explore correlates between sensory systems in honey bees and bumblebees (e.g., Erber et al., 2006; Marchal et al. 2029; Nouvian & Galizia 2020). Phototaxis has also been used to assess pesticide effects in honey bees (Bergougnoux et al., 2013; Charreton et al., 2015; Tosi and Nieh, 2017). These studies use vertical arenas and assess various endpoints related to locomotor function, such as ability to climb the arena, path trajectory, hyperactivity, and the time spent in defined areas across the light gradient.

Phototaxis is a fundamental component of bee life history, and has been associated to division of labour in social species. In honey bees, young nurse bees, which perform tasks inside of the hive, exhibit negative phototaxis, whereas older foragers exhibit positive phototaxis (Ben-Shahar et al., 2003). In bumblebees, in which division of labour is more closely associated to body size than age, nursing (mostly small) workers present lower levels of positive phototactic response than foraging (mostly large) workers (Merling et al., 2020). Solitary bees move towards light to emerge from their natal nest (Hirashima, 1972; Torchio, 1980; Szentgyörgyi & Woyciechowski, 2013), and nesting females also do so every time they leave their nest to go on a foraging trip. Because phototaxis is such a fundamental behaviour, intraspecific variation in phototactic response is expected to be low, thus increasing the robustness and repeatability of phototaxis-based methodologies. Our study has two objectives. First, to describe a simple phototaxis-based assay to assess sublethal pesticide effects in bees. Second, to validate the assay by demonstrating its ability to obtain ecotoxicological endpoints in both solitary bees and honey bees.

## 2. Methods and Materials

### 2.1 Phototaxis arena

We built a simple phototaxis box made of opaque corrugated plastic, measuring 50 x 50 x 85 cm (Fig. 1). An infra-red camera connected to a computer is fitted at the top of the box. There is a 1.5-cm-diameter hole at the bottom of one side of the box through which light from a LED bulb (1937.5 lux) is projected from outside. The hole is covered with white filter paper to homogenize the direction of the light and reduce brightness. The bee is introduced by lifting the box from its base. The bee is placed in a semi-circular start area at the base of the box and then it is covered with an opaque lid (e.g., the base of a Petri dish painted black) for acclimation. A string is attached to the lid so that it can be easily lifted to release the bee. The base of the box is lined with a sheet of white filter paper that is replaced after each individual trial to eliminate potential olfactory cues from previous tests.

**Fig. 1.**
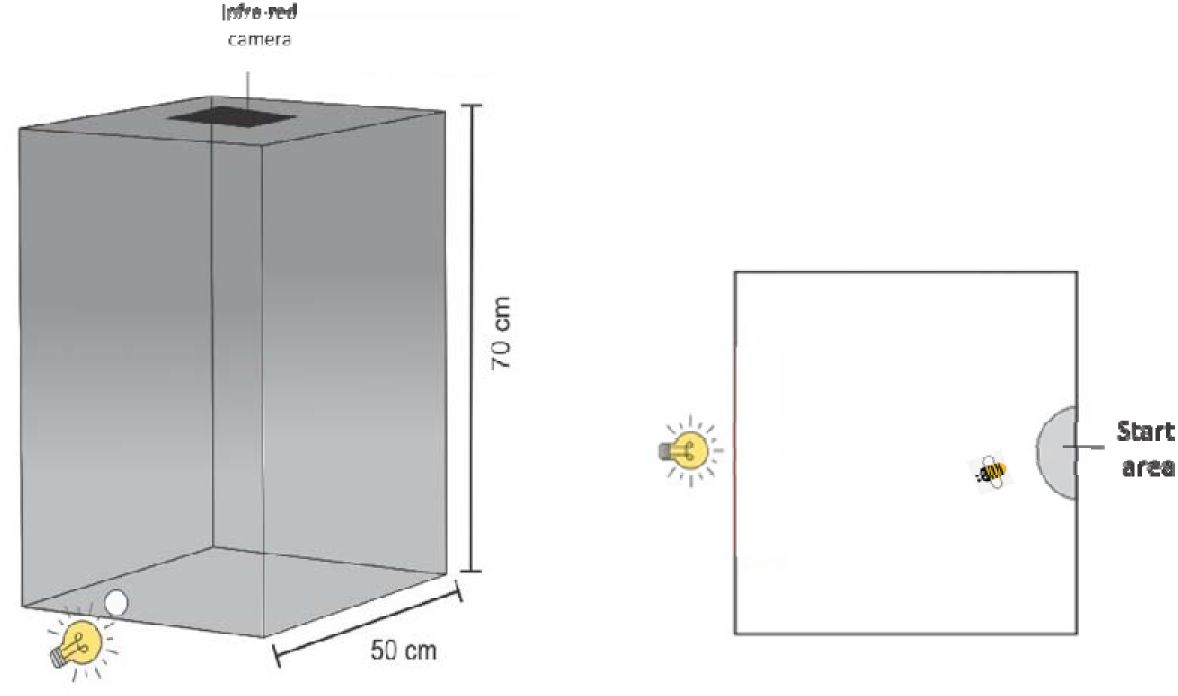
Phototaxis box. The bee is filmed with an infra-red camera as it walks across a horizontal plane (the floor of the box).

### 2.2 Test procedure

Bees in small cages (height: 7 cm; diameter: 10 cm) are orally exposed to the desired doses or concentrations of the test solution using the petal method (described in Azpiazu et al., 2023b). The exposed bee is left in the cage for two hours to allow the pesticide to take effect. The bee is then introduced into the phototaxis arena and covered with the opaque lid. Following an acclimation period of three minutes in total darkness, the light source is switched on, the lid is lifted and the infrared camera starts recording. Upon lifting the lid, the bee may be facing in any direction but this does not affect the outcome of the test because bees with a positive response quickly reorient towards the light. In any case, time is only measured from the moment the bee walks out of the start area.

Several parameters can be obtained from the video recordings, including: a) whether the bee reaches the light source within a certain time period; b) the time taken to reach the light source; c) the distance walked; d) the walking speed; e) various measures related to the bee’s trajectory, such as number and direction of turns; f) abnormal behaviours such as inactivity, walking backwards and trembling. However, in preliminary trials with bees exposed to various sublethal pesticide doses and with control (unexposed) bees of four species (*A. mellifera, O. cornuta, O. bicornis and Osmia tricornis*), we basically observed three distinct responses: 1) Bees that remained motionless in the start area for more than two minutes (no response); in our initial trials, this response was rare (2-5 %); 2) Bees that clearly walked towards the light source following a fairly straight trajectory (positive response) (<u>Video 1</u>); these bees typically reached the light source in less than one minute; all control bees exhibited this response; 3) Bees that walked erratically (neutral response) (<u>Video 2</u>); these bees walked around the arena and typically did not reach the light source even after extended observation periods (10-15min). We never observed bees attempting to fly or climbing the walls of the arena. Based on these three distinct responses, we decided to limit the duration of the test to 1 minute. This not only greatly reduces the time invested in each trial but also enabled us to work with a binary variable (positive response *vs* neutral response), thus simplifying data processing and analysis.

### 2.3 Validation of the test

To validate the phototaxis test, we conducted two experiments. In Experiment 1, we built dose-response curves for *O. bicornis* and *O. tricornis* females exposed to acetamiprid, a cyano-substituted neonicotinoid insecticide. These curves were used to calculate the median effective dose (ED50; the dose at which 50% of the individuals failed to respond positively to light) and the benchmark dose (BMD), the estimated lowest dose that produces an adverse response compared to the control. In Experiment 2, we measured the sublethal effects of an Ergosterol biosynthesis inhibitor (EBI) fungicide (tebuconazole) and three doses of a neonicotinoid insecticide (acetamiprid), both alone and in combination. To demonstrate that the assay could be used with various bee species, including both solitary and social ones, Experiment 2 was conducted with *O. cornuta* and *A. mellifera*. Figures S1 and S2 show the distribution of times taken to reach the light source by the four bee species used in the two experiments.

#### 2.3.1 Experiment 1: Dose-response curves

We worked with *O. bicornis* and *O. tricornis* populations reared at CREAF (Centre for Ecological Research and Forestry Applications). In April 2021, cocoons containing adults were removed from the wintering chamber (3-4 °C). We selected large cocoons to increase the likelihood of them containing females and to reduce variation in body size. Cocoons were incubated at 20 °C until emergence (Fig. 2). We worked with four batches of cocoons incubated over four consecutive weeks. On the day of peak emergence for each batch, newly-emerged females were collected and evenly distributed across treatments (see below). Bees were kept for 24 h in a screen cage (60 x 40 x 40 cm) for meconium deposition and fasting. Bees were then individually transferred to small cages (plastic containers with perforated lids; height: 7 cm; diameter: 10 cm) where they were orally exposed to 20 µl of a feeding solution (33% sucrose in distilled water; w/w) with a range of doses of Carnadine® (Acetamiprid 20%). Following preliminary trials, we orally exposed bees to six doses (*O. bicornis* : 0, 33.96, 52.63, 81.58, 126.45 and 196 ng/bee; *O. tricornis* : 0, 7.75, 12.01, 18.62, 28.86, 44.73 ng/bee) of a. i. in a geometric series of 1.55.

**Fig. 2.**
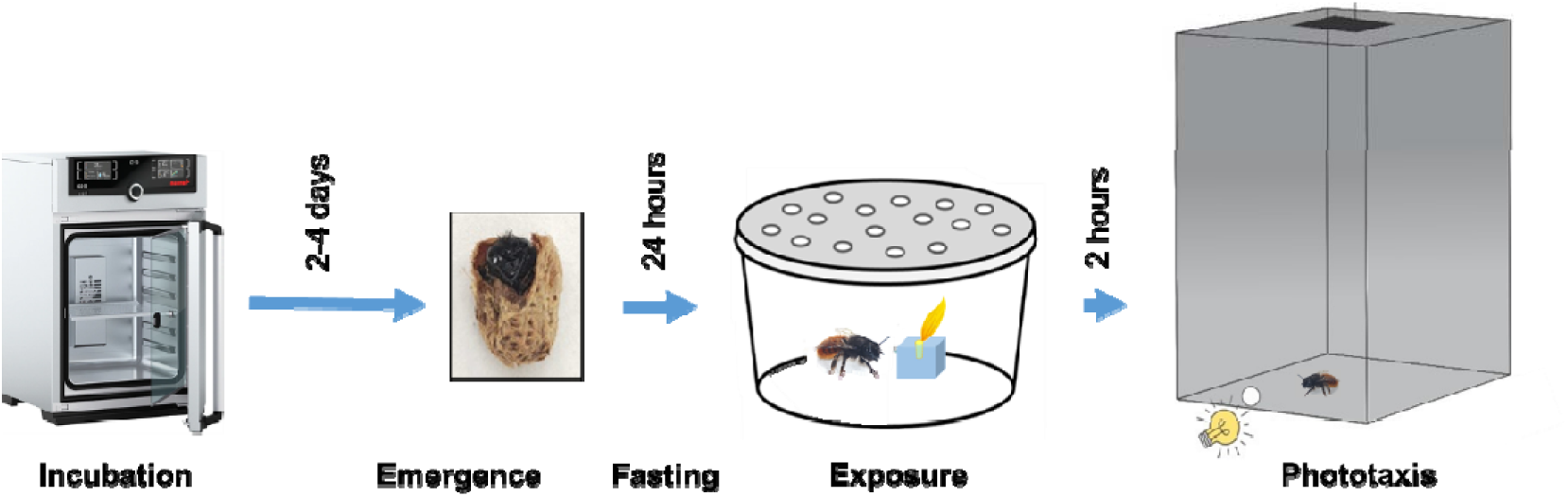
Experimental procedure used to test response to light (phototaxis) in newly-emerged *Osmia* spp. females orally exposed to sublethal pesticide doses.

Bees were exposed using the petal method (Azpiazu et al., 2023b). Feeding cages were checked every 15 minutes for 1.5 hours. After each check, bees that had consumed the entire solution were left in the feeding cages for 2 hours to allow the insecticide to take effect. Bees were then submitted to the phototaxis test. The entire procedure was conducted at room 22 °C). Bees that had consumed only part of the solution at the time of the check were discarded, as were those that had not fed at all after 1.5 hours. The procedure is summarized in Figure 2. Tested bees were returned to their individual cages and kept for 24 h to assess post-test mortality. Sample sizes were 30-31 bees per dose in *O. bicornis* and 20-23 in *O. tricornis*.

To rule out any potential effect of week on phototactic response, we ran preliminary generalized linear models (GLM with binomial error distribution and logit link function). Week and the interaction treatment x week were non-significant for both species (Table S1). Hence, week was not included in further analyses. Dose-response curves were estimated with the *drc* package (Ritz et al., 2015). The *mselect* function was used to determine which model was most appropriate for each species based on the Akaike Information Criterion (AIC). ED50s were estimated with the *ED* function of the *drc* package. To calculate the lowest levels of exposure at which our test could detect sublethal effects, we used a benchmark dose (BMD) approach. This is more reliable than the no-observed-averse-effect-level (NOAEL) approach because it is less dependent on dose selection and sample size (Davis et al., 2011). We established a 10% level of benchmark response (BMR) and used the dose-response data to estimate the BMD 95% lower confidence bound (BMDL) with the US EPA’s Benchmark Dose Software (BMDS Online version) (USEPA, 2023).

#### 2.3.2 Experiment 2: Pesticide mixtures

We worked with *O. cornuta* females from a population reared at CREAF and *A. mellifera* workers from a hive at the apiary of the School of Veterinary Science of the Autonomous University of Barcelona (UABEE). We followed the procedure described in Experiment 1 to incubate and handle newly-emerged *O. cornuta* females. Bees emerging on a given day were evenly distributed across treatments. *A. mellifera* workers were obtained by temporarily covering the entrance of the hive. Returning foragers attempting to enter the hive were gently picked up with a brush and placed in a glass jar with a perforated lid for ventilation. Using the petal method (C. Azpiazu et al., 2023b), *O. cornuta* females were individually and orally exposed to 20 µl of test solution (33% sucrose in distilled water, w/w, with the desired concentrations of fungicide and insecticide). Following OECD procedures (OECD, 1998), and because isolated honey bee workers are prone to die within hours, *A. mellifera* were exposed in groups of ten individuals to 200 µl of test solution. Under these conditions, all individuals reach similar levels of exposure through trophallaxis (food exchange) (OECD, 1998). We used Folicur**®** 25WG (tebuconazole 25%) and Carnadine**®** (Acetamiprid 20%). The fungicide dose (3000 ng/bee) was based on the Folicur**®** application rate (0.6 Kg/1000 l) recommended for apple orchards. We worked with three insecticide doses. The lowest dose (0.18 ng of a. i. /bee) is based on the concentration of acetamiprid found in oilseed rape nectar (7.6 ng/g; Pohorecka et al., 2012) and the intermediate dose (1 ng of a. i. /bee) on the concentration found in apple nectar (65 ng/g; Heller et al., 2020) (see calculations in Supplementary Information). The highest dose (2 ng of a.i./bee) is lower than the 100-fold dilution of the LD10 at 24 h of acetamiprid in *O. cornuta* (281.7 ng/bee; Barnadas, 2022). Two hours after exposure, bees were submitted to the phototaxis test. Sample sizes were 18-24 bees per species and treatment.

As in Experiment 1, we ran preliminary GLMs to rule out any potential effect of week on phototactic response. Week and the interaction treatment x week were again non-significant for both species (Table S2). We used a GLM to analyse the percentage of positive response by fitting a binomial error distribution with a clog-log link function. We included treatment, bee species and their interaction as fixed factors. We calculated *p* values of fixed effects using likelihood ratio tests. Pairwise comparisons were performed with Tukey’s p-value adjustment method (Lenth et al., 2019). Analyses were conducted in R (R Core Team., 2020).

## 3. Results

### 3.1 Experiment 1: Dose-response curves

Only 5.2 % of the *O. bicornis* and 2.4 % of the *O. tricornis* individuals failed to leave the start area two minutes after the lid was lifted. These bees were discarded. All control bees exhibited a positive response and reached the light source within one minute. The percentage of bees not reaching the light within one minute (neutral response) increased with dosage in both species (Fig. 3). ED50 was 102.26 ng/bee in *O. bicornis* and 22.79 ng/bee in *O. tricornis* (Table 1). Mortality at 24 h post-exposure was 0 in all treatments. Benchmark doses (BMD) corresponding to 10% failure to respond to light and their associated parameters are shown in Table 2. The lower confidence limits (BMDL) were 28.52 ng/bee in *O. bicornis* and 8.00 ng/bee in *O. tricornis*.

**Fig. 3.**
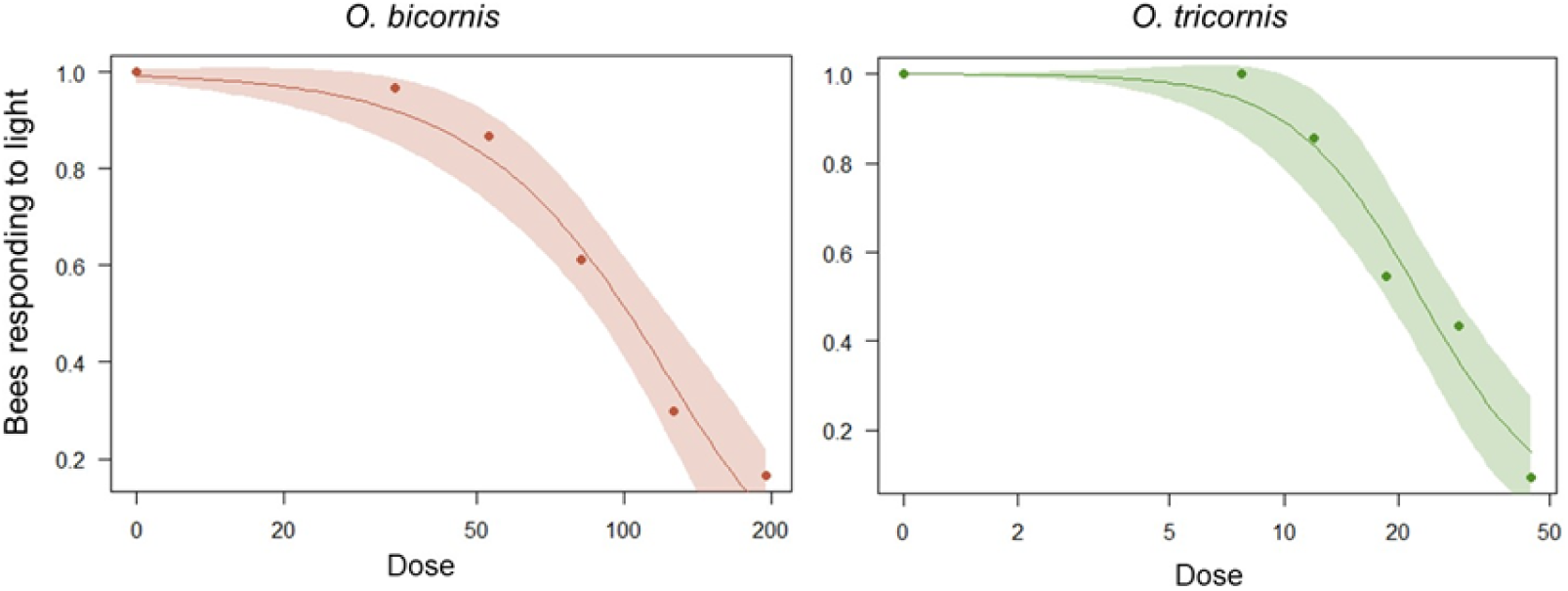
Proportion of *O. bicornis* and *O. tricornis* females responding positively to light after acute exposure to a range of doses (ng/bee) of acetamiprid. N= 30-31 (*O. bicornis*) and 20-23 (*O. tricornis*) females per dose.

**Table 1.** Median effect dose (ED50; dose at which 50% of bees tested failed to respond positively to light) and 95% confidence intervals of the insecticide acetamiprid in *O. bicornis* and *O. tricornis* females. P-values < 0.05 indicate no significant differences from reference (control) values.

| Species | n | Model | ED50<br>(ng/bee) | 95% CI | t-value | p-value |
| --- | --- | --- | --- | --- | --- | --- |
| *O. bicornis* | 181 | Weibull | 102.26 | 86.04 - 118.44 | 15.52 | <0.001 |
| *O. tricornis* | 122 | Log-logistic | 22.79 | 18.20 -27.39 | 13.77 | <0.001 |

**Table 2.**
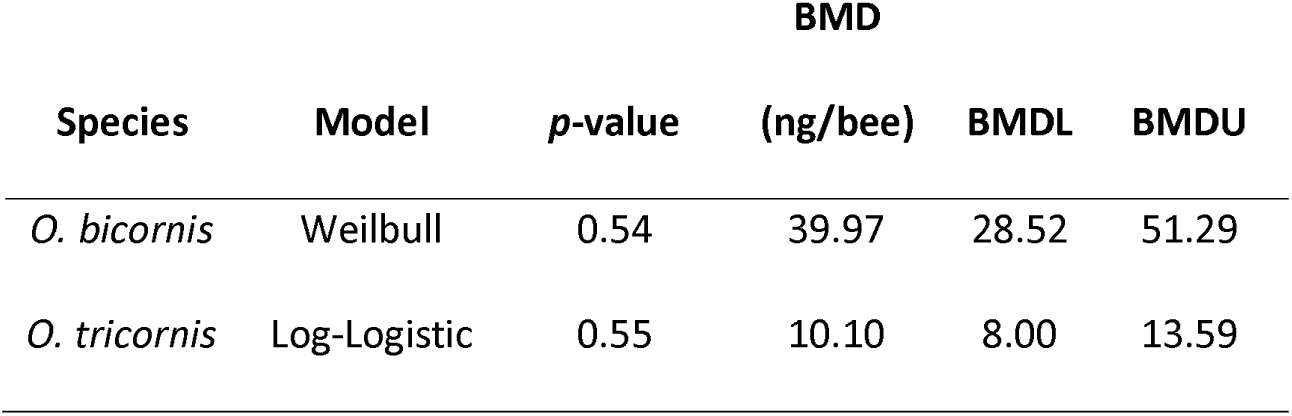
Benchmark dose (BMD) and its lower (BMDL) and upper (BMDU) confidence bounds of the insecticide acetamiprid eliciting a 10% failure to respond to light following oral acute exposure in *O. bicornis* and *O. tricornis.* P-values > 0.05 denote no significant differences between observed and predicted values.

### 3.2 Experiment 2: Pesticide mixtures

Only 14.1% (*O. cornuta*) and 3.2% (*A. mellifera*) of bees tested failed to respond to the test. These bees were discarded. Of the bees that did respond, all control individuals and all those exposed to the fungicide or any of the three doses of the insecticide alone responded positively to light (Fig. 4). However, the levels of positive response declined when bees were exposed to the mixture treatments (Fig. 4; Binomial GLM: treatment (χ^2^ = 88.4; df = 7; p < 0.0001); species (χ^2^ = 6.8; df = 1; p = 0.009); treatment x species (χ^2^ = 0.8; df = 7; p = 0.9).

**Figure 4.**
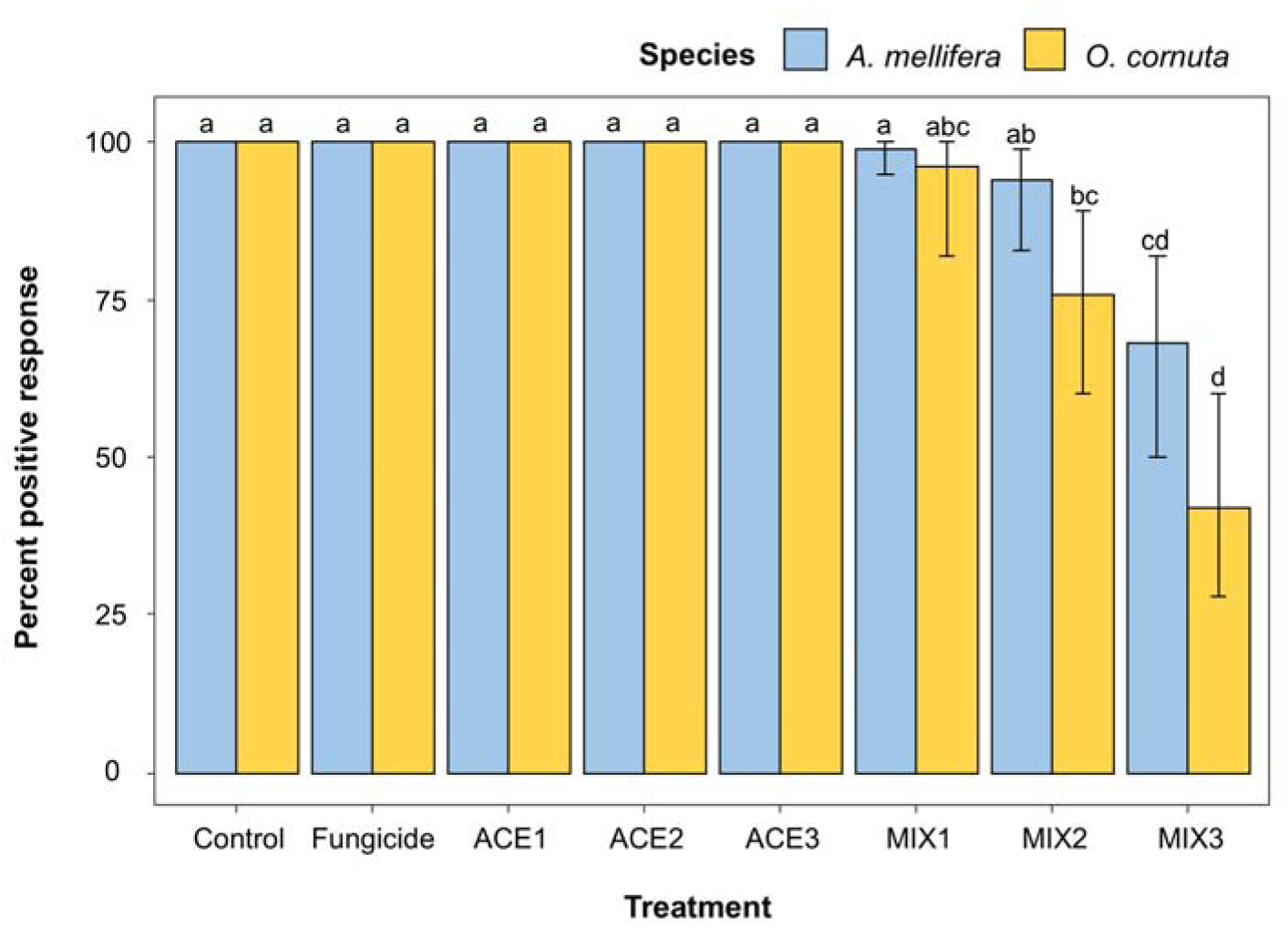
Model-estimated means and 95% confidence intervals for the percentage of positive responses to the phototaxis test in *O. cornuta* females and *A. mellifera* workers acutely exposed to the fungicide Folicur**®** (a.i., tebuconazole) and three doses of the insecticide Carnadine**®** (a.i., acetamiprid), alone and in combination. Different letters denote significant differences across treatments and species (Tukey HSD test, *p*lll<lll0.05). N= 18-24 bees per species and treatment.

## 4. Discussion

The objective of this study was to describe and validate an assay to measure the effects of sublethal doses of pesticides and combinations thereof on bees in the laboratory. We wanted our methodology to be simple (easy-to-implement), effective (a high proportion of individuals should respond to the test) and unambiguous (providing a clear, easily analysable response). The phototaxis assay described herein fulfils these three conditions. First, restricting the duration of the test to one minute, significantly reduces the time required to process each bee. In our laboratory, a team of two people could use two phototaxis boxes to process 50 bees per day, including the pesticide exposure and phototaxis phases. Second, the test is highly effective, even when bees are exposed to the higher insecticide concentrations. Of the 694 bees tested in our two experiments, only 6.63 % failed to respond, thus minimising the number of bees needed to reach a sufficient sample size. Third, our test provides an unambiguous binary response that does not require post-test elaboration. Importantly, the assay also enables measurement of other behavioural variables, such as the time taken to reach the light source, the length and trajectory of the path taken, and the walking speed, which could provide additional valuable endpoints for future studies.

Previous research (Bergougnoux et al., 2013; Charreton et al., 2015; Tosi and Nieh, 2017) assessed the effect of exposure to insecticides and other compounds on movement towards light in honey bees. These studies worked with vertical arenas and analysed endpoints related to locomotor function, such as path trajectory, impaired ability to ascend the arena, hyperactivity, and the time spent in areas with different levels of light intensity. These endpoints could also be measured with our assay, but the quick and dichotomous response of the bees tested rendered this unnecessary. Importantly, the erratic behaviour of bees responding neutrally to light was similar in solitary (*Osmia* spp.) and highly social bees (*A. mellifera*), indicating that the assay can be used across a range of bee species with highly contrasting life histories.

The mechanisms explaining the reduced response to light following insecticide exposure require further research. In *Drosophila melanogaster*, exposure to sublethal doses of the neonicotinoid insecticide imidacloprid was found to affect photoreceptor function, a proxy for blindness (Martelli et al., 2020). We recorded the behaviour of control bees in the phototaxis arena in total darkness. These bees typically remained motionless for a few seconds, and then started moving erratically in different directions for several minutes (<u>Video 3</u>). These bees often walked along the sides of the box (thigmotactic response) and sometimes tried to walk upwards (negative geotactic response), behaviours that could be interpreted as attempts to find an exit. In contrast, pesticide-exposed bees that did not respond positively to light typically started walking sooner and exhibited a greater tendency, at least initially, to walk toward the light source (<u>Video 4</u>). These observations suggest that impaired capacity to perceive light may not be the only or main mechanism explaining the behaviour of insecticide-exposed bees failing the phototaxis test. In another *D. melanogaster* study, individuals manipulated in various ways to impair their ability to fly exhibited reduced positive phototaxis (Gorostiza et al., 2016). In that study, normal levels of phototactic response were recovered once flying ability was restored, indicating that flies monitored their ability to fly and adjusted their photopreferences accordingly. A similar mechanism could explain the diminshed phototactic response in our study. Exposure to sublethal levels of insecticides could reduce the ability of bees to fly or otherwise cope with the environment, prompting their decision to ignore stimuli associated with the outside world.

We used the outcomes of the phototaxis test to build dose-response curves and calculate ED50 and BMDL values, thus demonstrating that the assay can be used to obtain relevant ecotoxicological endpoints in bee studies. In Experiment 1, the ED50 of acetamiprid was 102.26 ng/bee for *O. bicornis* and 22.79 ng/bee for *O. tricornis*. To our knowledge, oral LD50 data for acetamiprid are not available for these two species, but has been estimated at 893 ng/bee (at 24 h) for *O. cornuta* (Barnadas, 2022). Assuming a similar LD50 for *O. bicornis* and *O. tricornis*, the phototaxis test could detect median effects at doses that are around an order of magnitude lower than those of a lethal test. Comparison between the phototaxis and lethal toxicity tests can also be assessed with the sublethal toxicity ratio (SubTR) based on the lowest-observed-adverse-effect-level (LOAEL) (BMDL in our study) and the LD50 (Tosi et al., 2022). Again assuming an LD50 similar to that of *O. cornuta*, the SubTR in the tested *Osmia* species ranged between 0.03 and 0.009, indicating that the phototaxis response is a highly sensitive endpoint (Tosi et al., 2022).

Acetamiprid is one of the few neonicotinoid insecticides that are still authorized for outdoor use in the EU and is widely used on orchard and oilseed crops (EFSA, 2016). Due to its lower toxicity compared to other neonicotinoids (Pisa et al., 2015) it is sometimes advertised as “pollinator-friendly”. The results of Experiment 2 confirm its low toxicity at field realistic doses (0.18-2 ng per bee). However, when mixed with tebuconazole, doses of 1 and 2 ng/bee impaired the ability of *O. cornuta* and *A. mellifera*, respectively, to respond positively to light. Synergistic effects between neonicotinoid insecticides and EBI fungicides have been well documented (Iverson et al., 2019; Raimets et al., 2018; Robinson et al., 2017; Sgolastra et al., 2017) and are mediated by the inhibition of P450 detoxification systems (Schuhmann et al., 2022). Further validation of our assay stems from a study that assessed the combined effects of climate change and sublethal insecticide exposure in *O. cornuta* females (Albacete et al., 2023). In that study, bees were exposed to sublethal doses of sulfoxaflor, a sulfoximine insecticide that, like neonicotinoids, acts as a selective agonist of Nicotinic Acetyl Choline Receptors. The percentage of bees that responded positively to light decreased from 96 % in bees exposed to 0 ng of a. i. per bee, to 90 % in bees exposed to 4.55 ng of a. i. per bee, and to 54 % in bees exposed to 11.64 ng of a. i. per bee (Albacete et al., 2023), thus demonstrating that our assay also works with environmentally relevant exposure levels of another insecticide.

Sublethal effects are insufficiently addressed in current bee-pesticide risk assessment schemes, and the few tests available for honey bees are not easily applicable to other bee species (Sgolastra et al., 2019). Our study helps bridge this gap by providing a simple methodology that can detect sublethal effects at field realistic levels of exposure to acetamiprid as low as 1-30 ng/bee. Our methodology can be applied to both honey bees and solitary bees, thus allowing for direct comparison of intrinsic sublethal sensitivity across species with highly contrasting life histories. The validation of the assay yielded results that are congruent with previous ecotoxicology studies on bees at higher exposure levels, such as the higher sensitivity to neonicotinoids in *Osmia* spp. compared to *A. mellifera* (Arena and Sgolastra, 2014; Sgolastra et al., 2017; Uhl et al., 2019) and the higher toxicity of sulfoxaflor compared to acetamiprid (Lewis et al. 2016; Azpiazu et al., 2021; Barnadas, 2022). Our assay could be used to screen the effects of low (field realistic) levels of exposure across a range of bee species, thus enhancing our ability to protect this important group of pollinators.

## Supporting information

Highlights

## Acknowledgments

The authors are thankful to M. Barnadas, E. Serratosa, P. Soler and J. Benrezkallah for their technical assistance. We are also thankful to Dr Abdelaali El Had and Dr Gerardo Caja for access to the UABee apiary. Special thanks to Dr. Rentao Liu for his advice on journal publication.

## Author contributions

GS, SA, CA, AR and JB conceived the study. GS, SA and CA designed the methodology and collected the data. SA and CA analysed the data. GS and JB led the writing with assistance by FS and CA. All authors contributed critically to the drafts and gave final approval for publication.

---

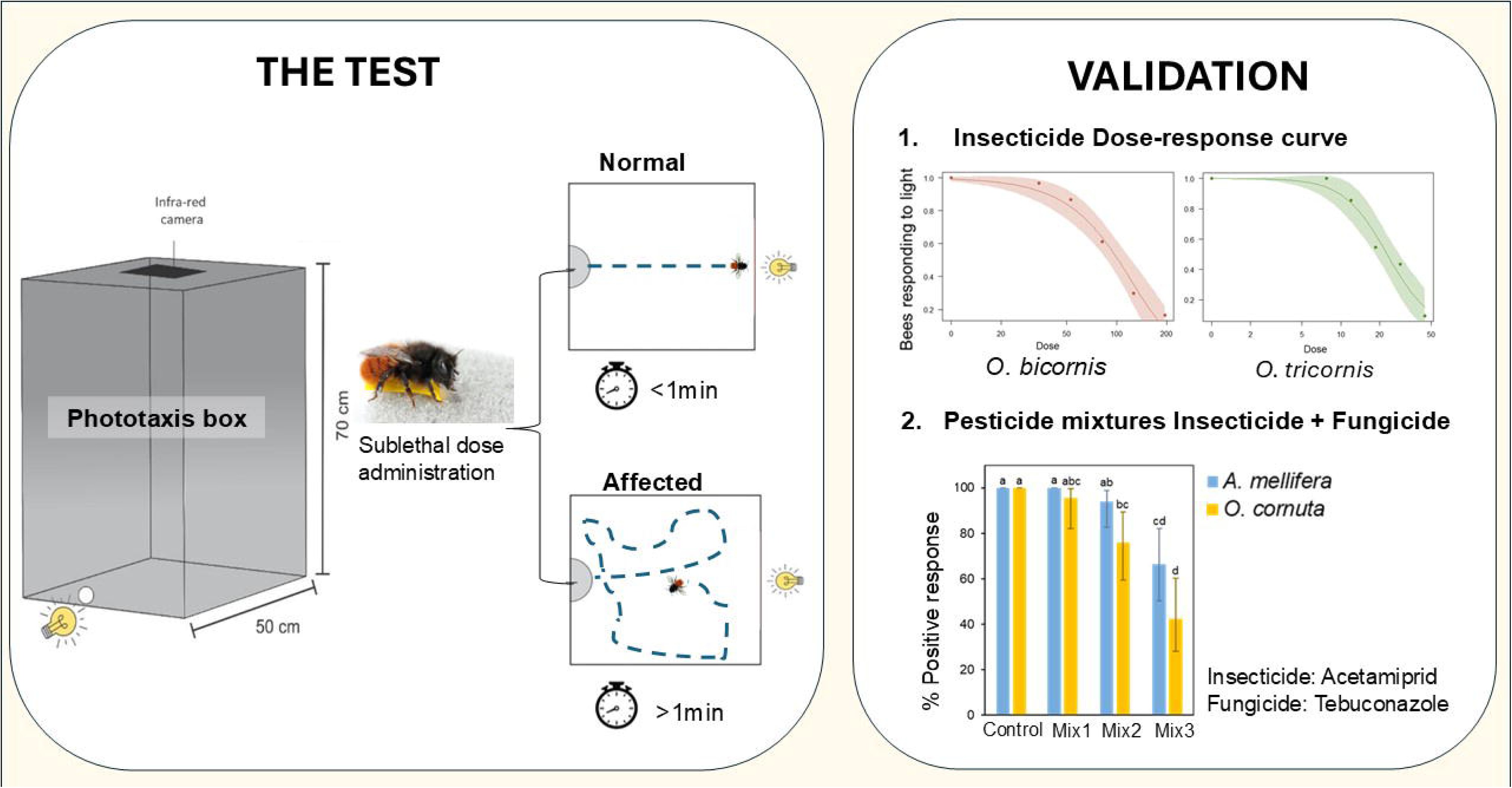

## References

Albacete, S., Sancho, G., Azpiazu, C., Rodrigo, A., Molowny-Horas, R., Sgolastra, F., Bosch, J., 2023. Bees exposed to climate change are more sensitive to pesticides. Glob. Chang. Biol. 29, 6248–6260. 10.1111/gcb.16928

Albacete, S., Sancho, G., Azpiazu, C., Rodrigo, A., Sgolastra, F., Bosch, J., 2024. Exposure to realistic levels of insecticide-fungicide mixtures affect reproductive success and population growth rates in a solitary bee. Environ. Int. 190 (2024) 108919. 10.1016/j.envint.2024.108919

Arena, M., Sgolastra, F., 2014. A meta-analysis comparing the sensitivity of bees to pesticides. Ecotoxicology 23, 324–334. 10.1007/s10646-014-1190-1

Azpiazu, C., Bosch, J., Bortolotti, L., Medrzycki, P., Teper, D., Molowny-Horas, R., Sgolastra, F., 2021. Toxicity of the insecticide sulfoxaflor alone and in combination with the fungicide fluxapyroxad in three bee species. Sci. Rep. 11, 6821. 10.1038/s41598-021-86036-1

Azpiazu, C., Medina, P., Sgolastra, F., Moreno-Delafuente, A., Viñuela, E., 2023a. Pesticide residues in nectar and pollen of melon crops: Risk to pollinators and effects of a specific pesticide mixture on Bombus terrestris (Hymenoptera: Apidae) micro-colonies. Environ. Pollut. 326, 121451. 10.1016/j.envpol.2023.121451

Azpiazu, C., Hinarejos, S., Sancho, G., Albacete, S., Sgolastra, F., Martins, C.A.H., Domene, X., Benrezkallah, J., Rodrigo, A., Arnan, X., Bosch, J., 2023b. Description and validation of an improved method to feed solitary bees (Osmia spp.) known amounts of pesticides. Ecotoxicol. Environ. Saf. 264, 115398. 10.1016/j.ecoenv.2023.115398

Barnadas, M., 2022. Toxicity of an insecticide-fungicide-adjuvant mixture in a solitary bee. MSc Thesis. Auton. Univ. Barcelona.

Ben-Shahar, Y., Leung, H.-T., Pak, W.L., Sokolowski, M.B., Robinson, G.E., 2003. cGMP-dependent changes in phototaxis: a possible role for the foraging gene in honey bee division of labor. J. Exp. Biol. 206, 2507–2515. 10.1242/jeb.00442

Bergougnoux, M., Treilhou, M., Armengaud, C., 2013. Exposure to thymol decreased phototactic behaviour in the honeybee (Apis mellifera) in laboratory conditions. Apidologie 44, 82–89. 10.1007/s13592-012-0158-5

Botías, C., David, A., Horwood, J., Abdul-Sada, A., Nicholls, E., Hill, E., Goulson, D., 2015. Neonicotinoid Residues in Wildflowers, a Potential Route of Chronic Exposure for Bees. Environ. Sci. Technol. 49, 12731–12740. 10.1021/acs.est.5b03459

Charreton, M., Decourtye, A., Henry, M., Rodet, G., Sandoz, J.-C., Charnet, P., Collet, C., 2015. A Locomotor Deficit Induced by Sublethal Doses of Pyrethroid and Neonicotinoid Insecticides in the Honeybee Apis mellifera. PLoS One 10, e0144879. 10.1371/journal.pone.0144879

Danforth, B.N., Minckley, R.L., Neff, J.L., 2019. The Solitary Bees: Biology, Evolution, Conservation. Princeton University Press. 10.2307/j.ctvd1c929

David, A., Botías, C., Abdul-Sada, A., Nicholls, E., Rotheray, E.L., Hill, E.M., Goulson, D., 2016. Widespread contamination of wildflower and bee-collected pollen with complex mixtures of neonicotinoids and fungicides commonly applied to crops. Environ. Int. 88, 169–178. 10.1016/j.envint.2015.12.011

Davis, J.A., Gift, J.S., Zhao, Q.J., 2011. Introduction to benchmark dose methods and U.S. EPA’s benchmark dose software (BMDS) version 2.1.1. Toxicol. Appl. Pharmacol. 254, 181–191. 10.1016/j.taap.2010.10.016

Decourtye, A., Armengaud, C., Renou, M., Devillers, J., Cluzeau, S., Gauthier, M., Pham-Delègue, M.-H., 2004. Imidacloprid impairs memory and brain metabolism in the honeybee (Apis mellifera L.). Pestic. Biochem. Physiol. 78, 83–92. 10.1016/j.pestbp.2003.10.001

Dicks, L. V., Breeze, T.D., Ngo, H.T., Senapathi, D., An, J., Aizen, M.A., Basu, P., Buchori, D., Galetto, L., Garibaldi, L.A., Gemmill-Herren, B., Howlett, B.G., Imperatriz-Fonseca, V.L., Johnson, S.D., Kovács-Hostyánszki, A., Kwon, Y.J., Lattorff, H.M.G., Lungharwo, T., Seymour, C.L., Vanbergen, A.J., Potts, S.G., 2021. A global-scale expert assessment of drivers and risks associated with pollinator decline. Nat. Ecol. Evol. 5, 1453–1461. 10.1038/s41559-021-01534-9

EFSA, 2016. Peer review of the pesticide risk assessment of the active substance acetamiprid. EFSA J. 14. 10.2903/j.efsa.2016.4610

EFSA, 2013. Guidance on the risk assessment of plant protection products on bees (Apis mellifera, Bombus spp. and solitary bees). EFSA J. 11. 10.2903/j.efsa.2013.3295

Erber, J., Hoormann, J., Scheiner, R., 2006. Phototactic behaviour correlates with gustatory responsiveness in honey bees (Apis mellifera L.). Behav. Brain Res. 174, 174–180. 10.1016/j.bbr.2006.07.023

Gong, Z., Tan, K., Nieh, J.C., 2019. Hornets possess long-lasting olfactory memories. J. Exp. Biol. 10.1242/jeb.200881

Gorostiza, E.A., Colomb, J., Brembs, B., 2016. A decision underlies phototaxis in an insect. Open Biol. 6, 160229. 10.1098/rsob.160229

Graham, K.K., Milbrath, M.O., Zhang, Y., Soehnlen, A., Baert, N., McArt, S., Isaacs, R., 2021. Identities, concentrations, and sources of pesticide exposure in pollen collected by managed bees during blueberry pollination. Sci. Rep. 11, 16857. 10.1038/s41598-021-96249-z

Hatjina, F., Papaefthimiou, C., Charistos, L., Dogaroglu, T., Bouga, M., Emmanouil, C., Arnold, G., 2013. Sublethal doses of imidacloprid decreased size of hypopharyngeal glands and respiratory rhythm of honeybees in vivo. Apidologie 44, 467–480. 10.1007/s13592-013-0199-4

Heller, S., Joshi, N.K., Chen, J., Rajotte, E.G., Mullin, C., Biddinger, D.J., 2020. Pollinator exposure to systemic insecticides and fungicides applied in the previous fall and pre-bloom period in apple orchards. Environ. Pollut. 265, 114589. 10.1016/j.envpol.2020.114589

Henry, M., Béguin, M., Requier, F., Rollin, O., Odoux, J.-F., Aupinel, P., Aptel, J., Tchamitchian, S., Decourtye, A., 2012. A Common Pesticide Decreases Foraging Success and Survival in Honey Bees. Science (80-.). 336, 348–350. 10.1126/science.1215039

Hirashima Y. (1972). Comparative studies on the orientation of pupae in the nests of solitary bees and wasps (Hymenoptera: Aculeata). Sci. Bull. Fac. Agr. Kyushu University 26 (1/4), 27–45.

Iverson, A., Hale, C., Richardson, L., Miller, O., McArt, S., 2019. Synergistic effects of three sterol biosynthesis inhibiting fungicides on the toxicity of a pyrethroid and neonicotinoid insecticide to bumble bees. Apidologie 50, 733–744. 10.1007/s13592-019-00681-0

Jékely G. 2009. Evolution of phototaxis. Phil. Trans. R. Soc. B3642795–2808. 10.1098/rstb.2009.0072

Jonason, D., Franzén, M., Ranius, T., 2014. Surveying Moths Using Light Traps: Effects of Weather and Time of Year. PLoS One 9, e92453. 10.1371/journal.pone.0092453

Kim, K., Huang, Q., Lei, C., 2019. Advances in insect phototaxis and application to pest management: a review. Pest Manag. Sci. 75, 3135–3143. 10.1002/ps.5536

Laloi, D., Sandoz, J.C., Picard-Nizou, A.L., Marchesi, A., Pouvreau, A., Taséi, J.N., Poppy, G., Pham-delègue, M.H., 1999. Olfactory conditioning of the proboscis extension in bumble bees. Entomol. Exp. Appl. 90, 123–129. 10.1046/j.1570-7458.1999.00430.x

Landim, C.D.C., 1967. Estudo comparativo de algumas glândulas das abelhas (Hymenoptera, Apoidea) e respectivas implicações evolutivas. Arq. Zool. 15, 177. 10.11606/issn.2176-7793.v15i3p177-290

Lenth, R., Singmann, H., Love, J., Buerkner, P., Herve, M., 2019. R Package ‘emmeans’. https://CRAN.R-project.org/package=emmeans.

Lewis, K. A., Tzilivakis, J., Warner, D. J. & Green, A. 2016. An international database for pesticide risk assessments and management. Hum. Ecol. Risk Assess. An Int. J. 22, 1050–1064.

Marchal P, Villar ME, Geng H, Arrufat P, Combe M, Viola H, Massou I, Giurfa M. Inhibitory learning of phototaxis by honeybees in a passive-avoidance task. Learn Mem. 2019 Sep 16;26(10):1–12. doi: 10.1101/lm.050120.119. PMID: 31527185; PMCID: PMC6749929.

Martelli, F., Zhongyuan, Z., Wang, J., Wong, C.-O., Karagas, N.E., Roessner, U., Rupasinghe, T., Venkatachalam, K., Perry, T., Bellen, H.J., Batterham, P., 2020. Low doses of the neonicotinoid insecticide imidacloprid induce ROS triggering neurological and metabolic impairments in Drosophila. Proc. Natl. Acad. Sci. 117, 25840–25850. 10.1073/pnas.2011828117

Mc Cabe, S.I., Hartfelder, K., Santana, W.C., Farina, W.M., 2007. Odor discrimination in classical conditioning of proboscis extension in two stingless bee species in comparison to Africanized honeybees. J. Comp. Physiol. A 193, 1089–1099. 10.1007/s00359-007-0260-8

Merling, M., Eisenmann, S., Bloch, G., 2020. Body size but not age influences phototaxis in bumble bee (Bombus terrestris, L.) workers. Apidologie 51, 763–776. 10.1007/s13592-020-00759-0

Nicholson, C.C., Knapp, J., Kiljanek, T., Albrecht, M., Chauzat, M.-P., Costa, C., De la Rúa, P., Klein, A.-M., Mänd, M., Potts, S.G., Schweiger, O., Bottero, I., Cini, E., de Miranda, J.R., Di Prisco, G., Dominik, C., Hodge, S., Kaunath, V., Knauer, A., Laurent, M., Martínez-López, V., Medrzycki, P., Pereira-Peixoto, M.H., Raimets, R., Schwarz, J.M., Senapathi, D., Tamburini, G., Brown, M.J.F., Stout, J.C., Rundlöf, M., 2023. Pesticide use negatively affects bumble bees across European landscapes. Nature. 10.1038/s41586-023-06773-3

Nouvian, M., Galizia, C.G. Complexity and plasticity in honey bee phototactic behaviour. Sci Rep 10, 7872 (2020). 10.1038/s41598-020-64782-y

OECD, 2021. Guidance document on Honey Bee (Apis mellifera L.) homing flight test, using single oral exposure to sublethaldoses of test chemical. Environ. Heal. Saf. Div.

OECD, 1998. Test No. 213: Honeybees, Acute Oral Toxicity Test. OECD Guidelines for the Testing of Chemicals, Section 2., OECD Guidelines for the Testing of Chemicals, Section 2. OECD. 10.1787/9789264070165-en

Pisa, L.W., Amaral-Rogers, V., Belzunces, L.P., Bonmatin, J.M., Downs, C.A., Goulson, D., Kreutzweiser, D.P., Krupke, C., Liess, M., McField, M., Morrissey, C.A., Noome, D.A., Settele, J., Simon-Delso, N., Stark, J.D., Van der Sluijs, J.P., Van Dyck, H., Wiemers, M., 2015. Effects of neonicotinoids and fipronil on non-target invertebrates. Environ. Sci. Pollut. Res. 22, 68–102. 10.1007/s11356-014-3471-x

Pohorecka, K., Skubida, P., Miszczak, A., Semkiw, P., Sikorski, P., Zagibajło, K., Teper, D., Kołtowski, Z., Skubida, M., Zdańska, D., Bober, A., 2012. Pozostałości Insektycydów Neonikotynoidowych w Nektarze i Pyłku Zbieranym Przez Pszczoły z Upraw Rzepaku i Ich Wpływ na Rodziny Pszczele. J. Apic. Sci. 56, 115–134. 10.2478/v10289-012-0029-3

Porrini, C., Mutinelli, F., Bortolotti, L., Granato, A., Laurenson, L., Roberts, K., Gallina, A., Silvester, N., Medrzycki, P., Renzi, T., Sgolastra, F., Lodesani, M., 2016. The Status of Honey Bee Health in Italy: Results from the Nationwide Bee Monitoring Network. PLoS One 11, e0155411. 10.1371/journal.pone.0155411

R Core Team., 2020. R: A language and environment for statistical computing. R Foundation for Statistical Computing. Available from https://www.r-project.org/.

Raimets, R., Karise, R., Mänd, M., Kaart, T., Ponting, S., Song, J., Cresswell, J.E., 2018. Synergistic interactions between a variety of insecticides and an ergosterol biosynthesis inhibitor fungicide in dietary exposures of bumble bees (<scp> *Bombus terrestris* </scp> L.). Pest Manag. Sci. 74, 541–546. 10.1002/ps.4756

Renzi, M.T., Rodríguez-Gasol, N., Medrzycki, P., Porrini, C., Martini, A., Burgio, G., Maini, S., Sgolastra, F., 2016. Combined effect of pollen quality and thiamethoxam on hypopharyngeal gland development and protein content in Apis mellifera. Apidologie 47, 779–788. 10.1007/s13592-016-0435-9

Ritz, C., Baty, F., Streibig, J.C., Gerhard, D., 2015. Dose-Response Analysis Using R. PLoS One 10, e0146021. 10.1371/journal.pone.0146021

Robinson, A., Hesketh, H., Lahive, E., Horton, A.A., Svendsen, C., Rortais, A., Dorne, J. Lou, Baas, J., Heard, M.S., Spurgeon, D.J., 2017. Comparing bee species responses to chemical mixtures: Common response patterns? PLoS One 12, e0176289. 10.1371/journal.pone.0176289

Russell Lenth, 2018. emmeans: Estimated Marginal Means, aka Least-Squares Means. R package version 1.3.1. https://CRAN.R-project.org/package=emmeans.

Schuhmann, A., Schmid, A.P., Manzer, S., Schulte, J., Scheiner, R., 2022. Interaction of Insecticides and Fungicides in Bees. Front. Insect Sci. 1. 10.3389/finsc.2021.808335

Sgolastra, F., Hinarejos, S., Pitts-Singer, T.L., Boyle, N.K., Joseph, T., Lūckmann, J., Raine, N.E., Singh, R., Williams, N.M., Bosch, J., 2019. Pesticide Exposure Assessment Paradigm for Solitary Bees. Environ. Entomol. 48, 22–35. 10.1093/ee/nvy105

Sgolastra, F., Medrzycki, P., Bortolotti, L., Renzi, M.T., Tosi, S., Bogo, G., Teper, D., Porrini, C., Molowny-Horas, R., Bosch, J., 2017. Synergistic mortality between a neonicotinoid insecticide and an ergosterol-biosynthesis-inhibiting fungicide in three bee species. Pest Manag. Sci. 73, 1236–1243. 10.1002/ps.4449

Siviter, H., Richman, S.K., Muth, F., 2021. Field-realistic neonicotinoid exposure has sub-lethal effects on non-Apis bees: A meta-analysis. Ecol. Lett. 24, 2586–2597. 10.1111/ele.13873

Szentgyörgyi H, Woyciechowski M. 2013. Cocoon orientation in the nests of red mason bees (Osmia bicornis) is affected by cocoon size and available space. Apidologie, 44, 334–341.

Takeda, K., 1961. Classical conditioned response in the honey bee. J. Insect Physiol. 6, 168–179. 10.1016/0022-1910(61)90060-9

Torchio, P.F. (1980) Factors affecting cocoon orientation in Osmia lignaria propinqua Cresson (Hymenoptera: Megachilidae). J. Kansas. Entomol. Soc. 53(2), 386–400.

Tosi, S., Nieh, J.C. A common neonicotinoid pesticide, thiamethoxam, alters honey bee activity, motor functions, and movement to light. Sci Rep 7, 15132 (2017). 10.1038/s41598-017-15308-6

Tosi, S., Sfeir, C., Carnesecchi, E., VanEngelsdorp, D., Chauzat, M.-P., 2022. Lethal, sublethal, and combined effects of pesticides on bees: A meta-analysis and new risk assessment tools. Sci. Total Environ. 844, 156857. 10.1016/j.scitotenv.2022.156857

Uhl, P., Awanbor, O., Schulz, R.S., Brühl, C.A., 2019. Is Osmia bicornis an adequate regulatory surrogate? Comparing its acute contact sensitivity to Apis mellifera. PLoS One 14, e0201081. 10.1371/journal.pone.0201081

USEPA, 2023. Benchmark Dose Tools (BMDS) Online [WWW Document]. URL https://bmdsonline.epa.gov.

USEPA, 2014. Guidance for Assessing Pesticide Risks to Bees. Off. Pestic. Programs.

Vorel, C.A., Pitts-Singer, T.L., 2010. The Proboscis Extension Reflex Not Elicited in Megachilid Bees. J. Kansas Entomol. Soc. 83, 80–83. 10.2317/0022-8567-83.1.80

Ward, L.T., Hladik, M.L., Guzman, A., Winsemius, S., Bautista, A., Kremen, C., Mills, N.J., 2022. Pesticide exposure of wild bees and honey bees foraging from field border flowers in intensively managed agriculture areas. Sci. Total Environ. 831, 154697. 10.1016/j.scitotenv.2022.154697

Zioga, E., Kelly, R., White, B., Stout, J.C., 2020. Plant protection product residues in plant pollen and nectar: A review of current knowledge. Environ. Res. 189, 109873. 10.1016/j.envres.2020.109873

