## Supplementary material for "A phototaxis assay to measure sublethal effects of pesticides on bees"

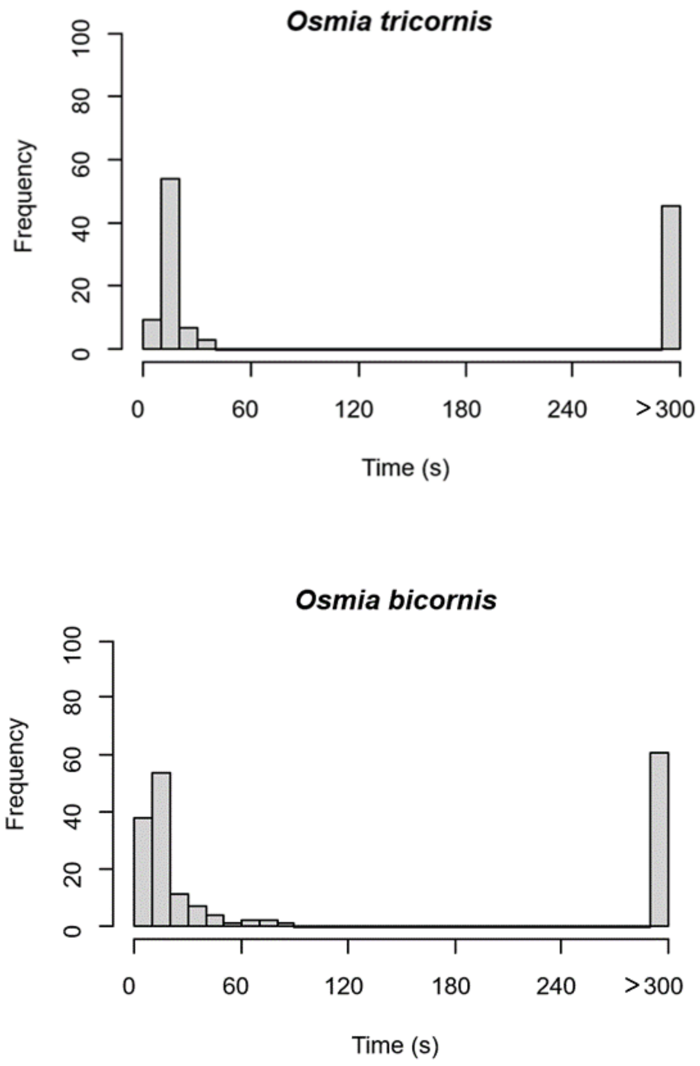


Fig. S1. Time to reach the light source in *O. tricornis* (n= 118) and *O. bicornis* (n= 181) females submitted to the phototaxis test in Experiment 1.

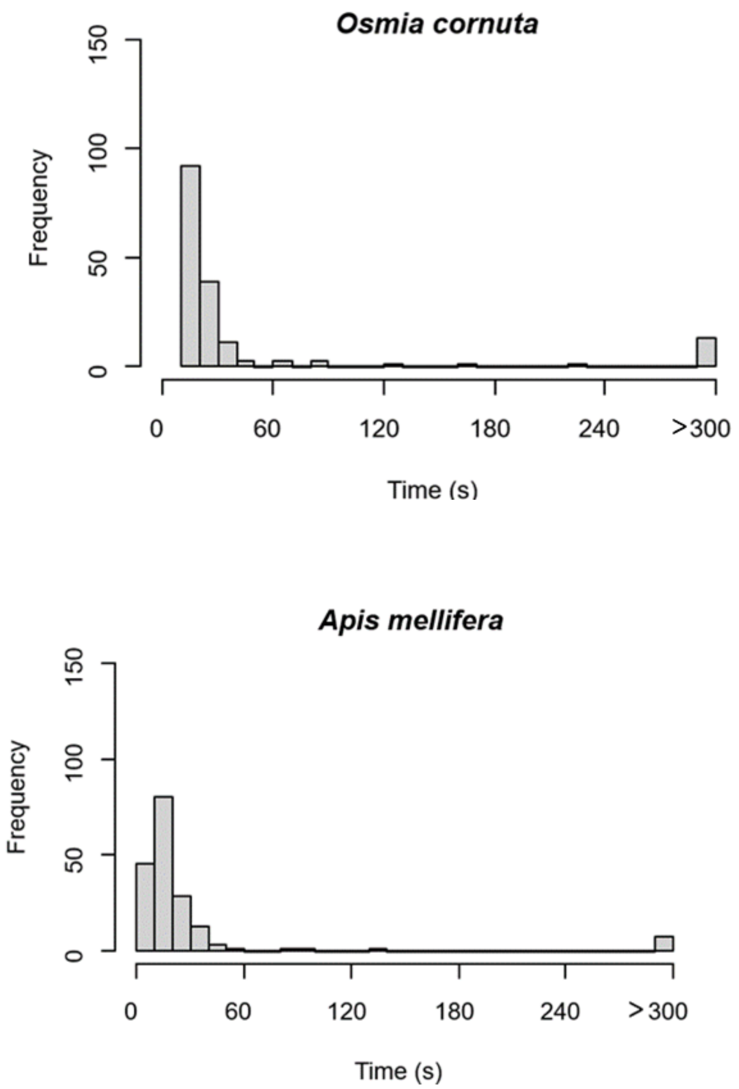


Fig. S2. Time to reach the light source in *O. cornuta* (n= 164) and *A. mellifera* (n= 181) females submitted to the phototaxis test in Experiment 2.

Calculations of Acetamiprid a. i. doses in experiment 2.

All bees were exposed to 20 µl of test solution (33% sucrose in distilled water, w/w) weighing 23 mg (density 1.15 g/mL).

**Dose 1.** Acetamiprid concentration reported by (Pohorecka et al. 2012) is 0.0076 ng of a. i. / mg of oilseed rape nectar. Therefore, the amount of a.i. per bee is 0.0076 ng/mg x 23 ng = 0.175 ng (0.18 ng per bee).

**Dose 2.** Acetamiprid concentration reported by (Heller et al. 2020) is 0.065 ng of a. i. / mg of apple nectar. Therefore, the amount of a.i. per bee would be 0.065 ng/mg × 23 mg = 1.49 ng. We decided to work with a lower (more conservative) concentration (1 ng/bee) so that our intermediate dose would be approximately half way between our lowest (0.18 ng/bee) and our highest (2 ng/bee) doses.

*Osmia tricornis*

| ***Model term*** | **$\chi^2$** | **df** | **p-value** |
|---|---|---|---|
| Pesticide treatment | 64.4 | 5 | **< 0.001** |
| Week | 1.4 | 3 | 0.705 |
| Pesticide treatment x Week | 13.1 | 15 | 0.772 |

*Osmia bicornis*

| ***Model term*** | **$\chi^2$** | **df** | **p-value** |
|---|---|---|---|
| Pesticide treatment | 107.7 | 5 | **< 0.001** |
| Week | 2.5 | 3 | 0.477 |
| Pesticide treatment x Week | 15.0 | 15 | 0.452 |

Table S1. Results of Binomial GLMs response ~ treatment * week, family = binomial (link = "logit" analysing the effect of treatment, week and their interaction on phototaxis response in *O. tricornis* and *O. bicornis* from Experiment 1.

*Osmia cornuta*

| ***Model term*** | **$\chi^2$** | **df** | **p-value** |
|---|---|---|---|
| Pesticide treatment | 56.9 | 7 | **< 0.001** |
| Week | 1.0 | 3 | 0.804 |
| Pesticide treatment x Week | 9.7 | 21 | 0.982 |

*Apis mellifera*

| ***Model term*** | **$\chi^2$** | **df** | **p-value** |
|---|---|---|---|
| Pesticide treatment | 38.3 | 7 | **< 0.001** |
| Week | 1.9 | 3 | 0.578 |
| Pesticide treatment x Week | 4.5 | 21 | 0.999 |

Table S2. Results of Binomial GLMs response ~ treatment * week, family = binomial (link = "logit" analysing the effect of treatment, week and their interaction on phototaxis response in *O. cornuta* and *A. mellifera* from Experiment 2.
