## Supplementary material for "A phototaxis assay to measure sublethal effects of pesticides on bees": Highlights

- Bee-pesticide risk assessment should address sublethal effects
- A phototaxis-based assay to measure sublethal effects is provided
- The assay works with both honeybees and solitary bees
- The assay detects effects at low field-realistic exposure levels of acetamiprid
- To validate the assay, we calculate dose-response curves and ED50s

Corresponding author:
Gonzalo Sancho:
